# Emergence of hierarchical organization in memory for random material

**DOI:** 10.1101/541490

**Authors:** Michelangelo Naim, Mikhail Katkov, Stefano Recanatesi, Misha Tsodyks

**Author notes:** These authors contributed equally to this work (Michelangelo Naim, Mikhail Katkov, Stefano Recanatesi).

## Abstract

Structured information is easier to remember and recall than random one. In real life, information exhibits multi-level hierarchical organization, such as clauses, sentences, episodes and narratives in language. Here we show that multi-level grouping emerges even when participants perform memory recall experiments with random sets of words. To quantitatively probe brain mechanisms involved in memory structuring, we consider an experimental protocol where participants perform ‘final free recall’ (FFR) of several random lists of words each of which was first presented and recalled individually. We observe a hierarchy of grouping organizations of FFR, most notably many participants sequentially recalled relatively long chunks of words from each list before recalling words from another list. Moreover, participants who exhibited strongest organization during FFR achieved highest levels of performance. Based on these results, we develop a hierarchical model of memory recall that is broadly compatible with our findings. Our study shows how highly controlled memory experiments with random and meaningless material, when combined with simple models, can be used to quantitatively probe the way meaningful information can efficiently be organized and processed in the brain, so to be easily retrieved.

**Significance Statement:** Information that people communicate to each other is highly structured. For example, a story contains meaningful elements of various degrees of complexity (clauses, sentences, episodes etc). Recalling a story, we are chiefly concerned with these meaningful elements and not its exact wording. Here we show that people introduce structure even when recalling random lists of words, by grouping the words into ‘chunks’ of various sizes. Doing so improves their performance. The so formed chunks closely correspond in size to story elements described above. This suggests that our memory is trained to create a structure that resembles the one it typically deals with in real life, and that using random material like word lists can be used to quantitatively probe these memory mechanisms.

## Introduction

One of the goals of neuroscience is to establish a link between neuronal processes and cognitive functions, such as memory, emotions, or language (1). Within this scope is memory recall, which is a challenging memory task for human participants, has been found to shed light on a number of memory and linguistic phenomena. For example, when retrieving random lists of words, people remember a very limited number of words even for lists of moderate lengths (2, 3). We showed recently that two simple principles are sufficient to explain free recall performance for randomly assembled lists of words for a wide range of list lengths (1):

1. The encoding principle states that each memory item is encoded (“represented”) in the brain by a specific group of neurons in a dedicated memory network. When an item is retrieved (“recalled”), either spontaneously or when triggered by an external cue, this specific group of neurons is activated.
2. The associativity principle for which, in the absence of specific retrieval cues, the currently retrieved item plays the role of an internal cue that triggers the retrieval of the next item.

From these two principles, we were able to theoretically derive that, on average, 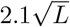 words out of the presented *L* words would be recalled. This simple expression is compatible with earlier experimental results (4) and with the very limited recall performance of individual lists of words (5), and was also confirmed by our recent experiments (6).

Nevertheless, people exhibit a striking ability to give long talks or recite lengthy poems by heart. How do they achieve this? And why are some people better than others in terms of their capacity to retrieve information from memory? It is well known that the more information is structured the easier it is to remember. For example, a story may contain distinct episodes that relate to each other in multiple logical chains that give rise to its ‘meaning’ and makes it more memorable (7–9). Often speakers introduce organization to the material to be communicated in order to improve its retention; one of the most prominent organizational strategies is grouping (the parceling of information into smaller parts). Pausing at appropriate places while speaking, allows listeners to divide the speech into meaningful parts (3). Evidence shows that encoding our experiences into distinct entities (‘chunks’), results in a hierarchical representation of information, that facilitates its subsequent retrieval (7, 10, 11). In support of this view, we recently showed that participants employing chunking as a retrieval strategy outperform other participants even when recalling random lists of words (12). In this study we demonstrate that in a memory task where information contains a limited degree of structure, best performing participants employ a hierarchical recall strategy. By means of a theoretical model we determine how properties of hierarchical representations, employing both representations of words and lists of words, account for most of the experimental observations and across subject variability.

## Grouping over lists, induced by presentation protocol

The protocol of the experimental dataset we analyze, obtained in the lab of Prof. Kahana at the University of Pennsylvania (13), adheres to the following structure. Each participant performed 16 Immediate Free Recall (IFR) trials a day with randomly assembled non-overlapping lists of 16 words. On selected days they were subsequently asked to recall all the words presented on that day (FFR; Fig. 1). Averaged over roughly 900 FFR sessions, participants recalled 57 words per session. This level of performance can be compared to the theoretical prediction resulting from modeling the experiment (14) according to which, out of a list of *L* random words, on average no more than 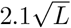 words would be recalled (1, 12). If one considers a FFR trial as the recall of a random list of *L* = 256 words, participants would on average recall 32 words, i.e. the observed performance of 57 words per session is almost twice the theoretical prediction. This discrepancy could be an indication that participants take advantage of the structural organization of presented words imposed by prior IFR trials. To prove that this is indeed the case, we quantify the level of grouping in FFR over the presented lists with quantitative measure for which each FFR trial is characterized by a value *p*_16_ that reflects the tendency to recall subsequent words from the same list before switching to another list (see Materials and Methods (12)). The distribution of *p*_16_ over the data is very wide Fig. 1b, covering the range from 0 (random recall) to 0.9 (strong degree of grouping; see Figs. 1c to 1e for three prototypical examples). Displaying the FFR performance versus the grouping measure *p*_16_ revealed a striking correlation between the two (*r* = 0.62, *p* = 4 × 10^-97^), with the bulk of data well characterized by linear dependence of performance on *p*_16_. Interestingly, in the limit *p*_16_ → 0, i.e. when no grouping is employed, performance approached a value of 30 words, supporting the theoretical prediction (1). We also observe that in FFR sessions with highest values of *p*_16_ participants occasionally recalled single words from a list in between longer sequences from other lists (Fig. 1c; see e.g. a single word from the 15th list recalled between two groups of words from the 4th list). We speculate that these short ‘intrusions’ are analogous to famous ‘slips of the tongue’ in natural speech (5).

**Fig. 1.**
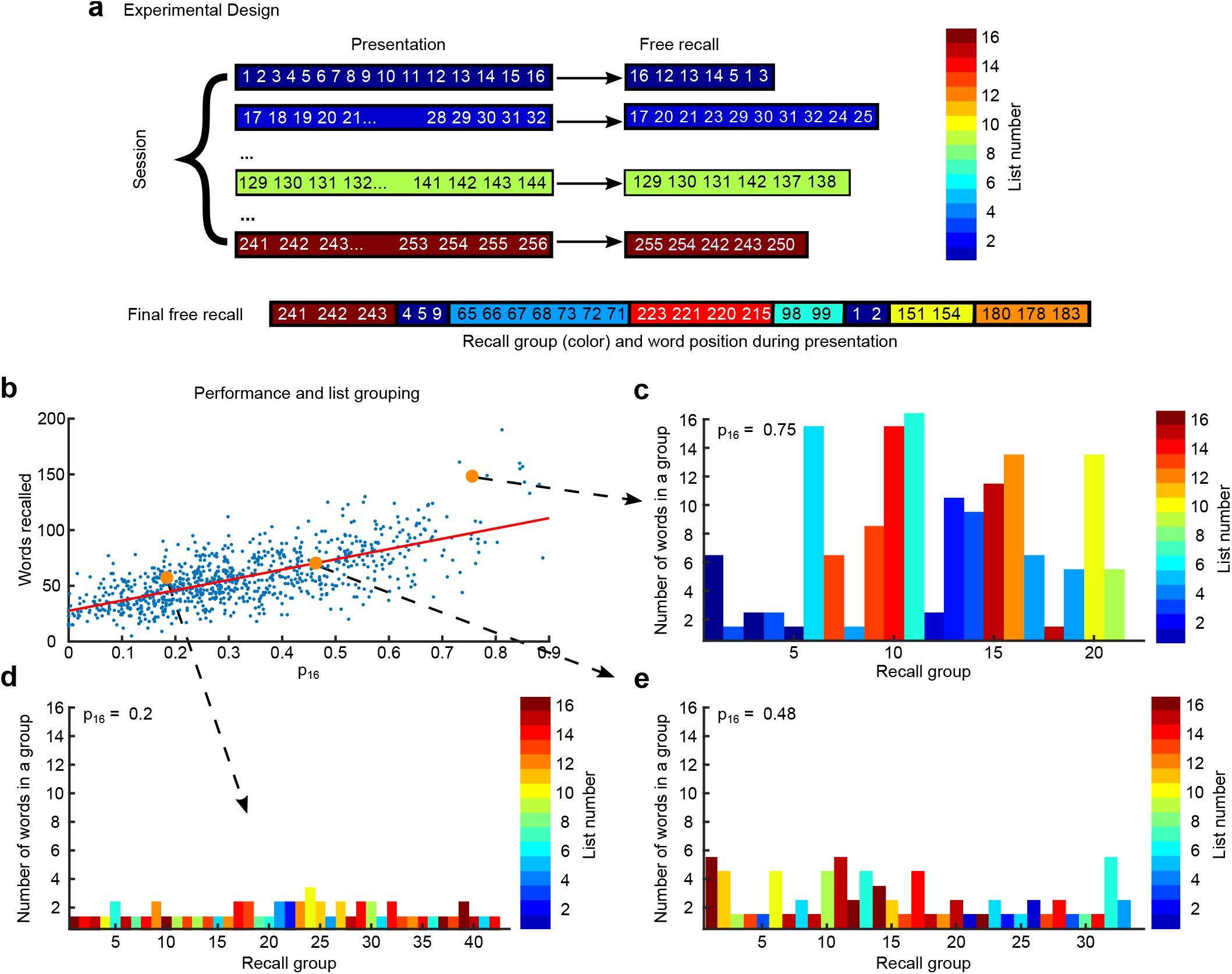
Experimental design and list-grouping results. **(a)** Experimental design – each day 16 lists were presented to human participants (colored lines on the left, with numbers representing serial presented position of the word during the day, and color representing the list number from blue to brown with color code presented in the right), after presentation of a list participants recalled as many words as they could (colored lines in the middle with a serial position of recalled words); at the end of some days participants performed final free recall (FFR), where they recalled as many words presented during the day as they could (bottom line with each recalled word colored according to the presentation list). **(b)** Number of recalled words during FFR vs grouping measure *p*16 (see details in the text); red line denotes linear fit. **(c-e)** are examples of FFR trials for three levels of grouping: high **(c)**, low **(d)** and intermediate **(e)**. All words consecutively recalled from the presented list are shown as a vertical column with color corresponding to list number and height to the number of words in the sequence. High level of grouping **(c)** is characterized by consecutive recalls of many words from the same list, sometimes interleaved with 1 or 2 words from other lists. Low level of grouping **(d)** is characterized by frequent switches between lists with 1 or 2 words recalled consecutively from the same list. Intermediate level of grouping **(e)** shows a mixture of high and low level grouping.

A possible interpretation of the above results is that participants perform FFR by applying a mixture of two recall strategies, one that treats all the words as one long random list, and another one that operates on two levels, namely individually presented lists and words within a list. As the second strategy gains prominence, recall becomes progressively more grouped and the value of *p*_16_ increases, accompanied by the increase in performance. In particular, the participants could develop stable representations of each list as a separate entity and ‘recall’ a list before recalling words from that list.

## Spontaneous grouping within presented lists

The grouping over lists exhibited in Fig. 1 is induced by the experimental protocol as lists are first presented and recalled individually in the IFR protocol. Another level of grouping, that was not induced by the protocol, was identified in FFR through the analysis of IFR data: a small proportion of participants develops chunking strategies in IFR (15, 16). These participants divide lists of 16 words into groups of 3 or 4 consecutively presented words and recall these chunks as single entities (12). This kind of chunking is not imposed by the protocol; hence, it must emerge from active manipulation of the presented list, for example representing chunks of words as separate items in memory. Here we wondered whether the chunks observed during IFR remained in memory till FFR trials. It is hard to infer whether chunking occurred in every single trial, hence we assumed that a chunk is recalled as a unit when all words from that chunk are recalled consecutively (not necessarily in the correct order). We therefore isolated all chunks of size 4 that were recalled during IFR trials, and considered the recall of the constituting words during FFR. We computed the probability for the different number of words from this chunk to be recalled. The results are shown in Fig. 2a. We found that for the first three chunks in the list, probability has two peaks, at 0 and 4 words, indicating the tendency for all 4 words in these chunks to be recalled or omitted as a single unit. Interestingly, the probability curve for the last chunk in a list decays monotonically, indicating that words from that chunk are recalled independently. A plausible explanation of this effect is that the last several words in a list are typically recalled immediately during IFR since they are maintained in working memory after the list is presented, and hence their recall is effortless and does not lead to the formation of a chunk representation in memory. A similar explanation also accounts for a recently reported ‘anti-recency’ effect in FFR, where the last words in a list have *lower* probability to be recalled, as opposed to the well-documented positive recency effect during IFR (17). For comparison, if the same analysis is performed for IFR trials where the same four words were recalled but with at least one intervening word, the corresponding probabilities do not exhibit a peak at four words recalled (Fig. 2b).

**Fig. 2.**
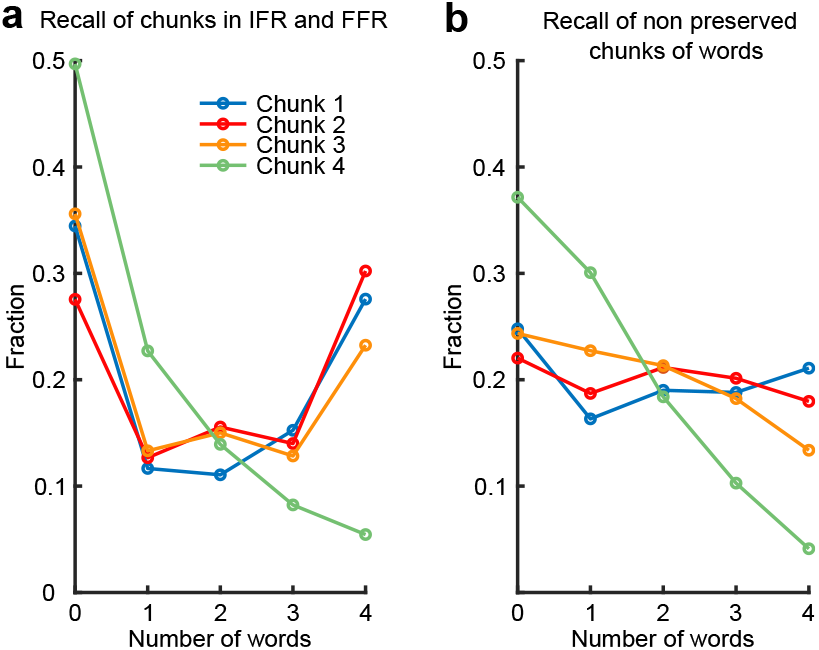
Fraction of chunk pieces of different length (number of words) recalled during final free recall (FFR). **(a)** When chunk is recalled as a whole (all 4 words from the chunk are recalled consecutively) in Immediate Free Recall (IFR). **(b)** When all words from a chunk recalled in IFR, but there are interleaving words from different chunks, i.e. a chunk is not recalled as a whole.

## Spontaneous grouping of lists

Some of the best participants who employ a strong over the list grouping imposed by the presentation protocol, also exhibit a higher-level grouping of lists. In particular, they tend to recall lists in chunks of four consecutive lists, as illustrated in figure Fig. 3.

**Fig. 3.**
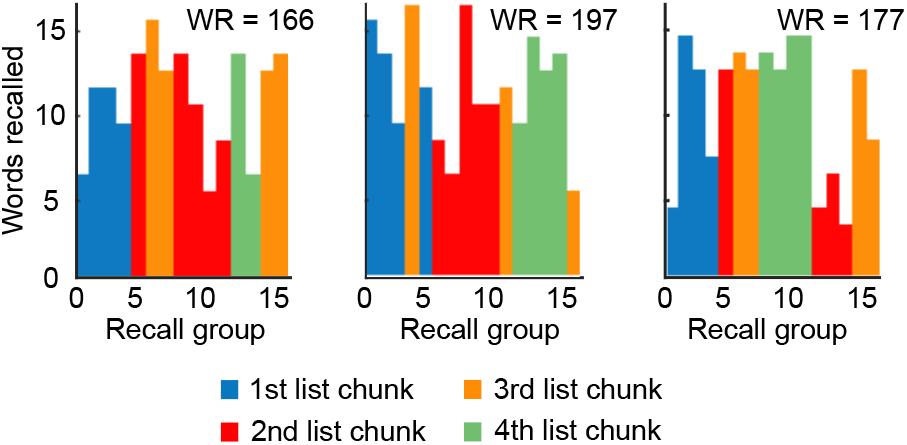
Three examples of FFR sessions with strong list chunking. Each plot displays the structure of a FFR session. Each bar represents a sequence of words recalled from a single list with the height of the bar indicating the number of words in a sequence and the color corresponding to the list chunk number. On top is reported the number of words recalled (WR) in each of these sessions.

Taking together, the results presented above illustrate that our memory is trained to create a structure on different levels of organization, including those that are not directly imposed by the presentation protocol.

## Hierarchical model of memory recall

In this section we show how the two principles presented above (encoding and associativity principle) account for the bulk of experimental findings. We build a hierarchical model of memory based on these two principles and show that it‘s behavior is in agreement with experimental results presented above.

### A. Modeling the Encoding

We extend the encoding principle formulated above for the recall of single lists in the following way. We postulate that different distinct levels of information (words, chunks, lists, context..) are encoded in the form of sparse random neuronal populations in the corresponding distinct subnetworks (see Fig. 4a), in line with the TCM model (18)). In the experimental paradigm words are presented in lists of 16 items and each session consists of 16 lists. Accordingly, each word *W* ls labeled by the triple of indexes: *W* = (*w, l, s*), corresponding to the presentation position in the list, the serial position of the list, and the session, respectively. Mathematically, each word *W* is represented by a binary pattern ***ξ***^*W*^ consisting of three parts:

**Fig. 4.**
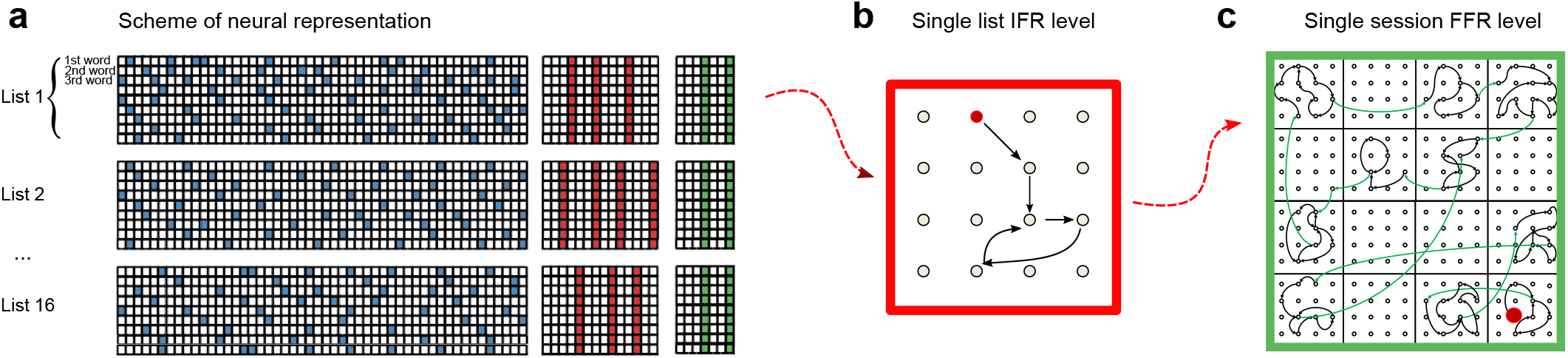
**(a)** Cartoon of the encoding of information in the dedicated memory networks. 3 subnetworks encoding words, lists and sessions are shown. Each row corresponds to a word and each column to a neuron. Colored neurons are encoding word (blue color), list (red color) or session (green color). Each word in word subnetwork is encoded by small fraction of random neurons, whereas words recalled in the IFR from the same list have the same encoding neurons in the list network. All words recalled in IFR from the same session have the same session encoding neurons. **(b)** An example of possible transitions induced by the model during the retrieval of 16 words from the same list. **(c)** An example of possible transitions in the FFR trial with large biding constant *E* (see details in text). Intralist transitions are colored in black, and extralist transitions are colored in green. Red circle is an initial item recalled.

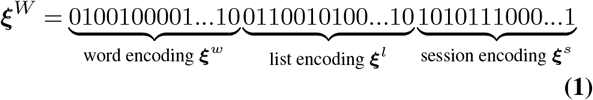

The length of the three vectors equals the number of neurons in each subnetwork *N*_*w*_, *N*_*l*_, *N*_*s*_. Each neuron encodes a word *W* with probability *f* so that the total number of neurons which encode a word *W* is on average *f* · (*N*_*w*_ + *N*_*l*_ + *N*_*s*_) = *f* · *N*.

In our previous studies (1, 19), transitions between words were driven by similarity matrices encapsulating resemblance of words representations and a neural dynamic inducing such transitions was proposed in (20, 21). The similarity between any two words *W*_1_ and *W*_2_ can be decomposed in three parts:

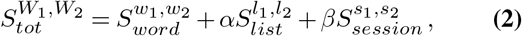

where 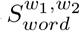 is the similarity matrix between words *W*_1_ and *W*_2_ in words subnetwork; 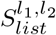 is the similarity between lists *l*_1_ and *l*_2_ in list subnetwork; 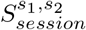 is the similarity of sessions *s*_1_ and *s*_2_ in the session subnetwork; parameters *α* and *β* reweigh the relative strength of the list and session context populations respectively in driving the retrieval process, in order to account for inter-subject variability (see below; cfr. Methods for details). The values of such similarities across all words determine the properties of the FFR process as we will further illustrate in modeling the associative principle and with it the retrieval dynamics.

A critical feature of the experimental dataset arises from the fact that words were not only presented in lists of 16 items, but also retrieved in immediate free recall IFR trials, Fig. 4b. The effect of IFR on the encoding process is crucial as 87% of the words recalled in IFR are later retrieved in FFR, as opposed to 13% of words not recalled in IFR. Because of the importance of IFR retrievals we model this phenomena by considering IFR as overruling the list presentation. In other words the similarity between two presented words induced by the list subnetwork will only be kept if both of the words were recalled in the subsequent IFR trial and otherwise will be put to zero (cfr. Methods).

### B. Associative transitions

The model of the encoding principle provides a simple mathematical characterization of words representation, but it does not describe how these representations are exploited in the retrieval dynamics. This is described within the scope of the associative principles which determines the rules that dictates transitions between words during the recall session.

According to the associativity principle the currently retrieved item functions as an internal cue that triggers the retrieval of the next one. This suggests that transitions between words are brought about by similarities between the active word – the last retrieved one – and other encoded words. Mathematically this means that if we assume the FFR session to start with any word *W*_1_, recalled during IFR, then transition to the next word *W*_2_ is driven by the total similarity 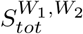. The word which is most similar to the one active is then activated and the process continues leading to the retrieval of more and more words. Importantly the last retrieved word cannot be activated so that a transition which just happened cannot immediately happen in the reverse direction. The dynamical recall process is solely driven by the associative principle and continues until it enters a loop when the same transition is repeated twice (see (14); cfr. Fig. 4b) after which the process would recapitulate the same words already retrieved: here is where the hierarchical representation of words comes at hand. When the process enters a loop, we assume that the next transition is overruled by a transition that does not follow the full similarity matrix *S*_*tot*_ to determine the most similar word to the last retrieved one, but rather:

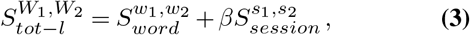

where the similarity between words arising from being in the same IFR trial *l* is suppressed. We call these transitions *random transitions* in contrast with the *structured transitions* induced by *S*_*tot*_. Upon triggering the retrieval of a new word through this transition the process reverts to using the full similarity matrix *S*_*tot*_ until it eventually enters a big loop Fig. 4c exemplifies this dynamic, the process starts in the word represented by the red dot, then a few transitions induced by *S*_*tot*_ take place (black arrows) until an already retrieved word is activated and a random transition occurs (green arrow). Then the same process starts over again until a random transition triggers the activation of a word already retrieved (blue arrow) which concludes the retrieval. Importantly, structured transitions are not confined to happen between words of the same list and similarly random transitions may happen between words that were in the same list or IFR trial.

The process just illustrated is mathematically formalized as a mixture model where transitions are triggered in either of two ways - structured transitions and random transitions of encoded word (cfr. Fig. 4c).

The hierarchical representation is exploited as, through it, different kind of transitions may occur. We will give reason for the biological plausibility of this model in the discussion, a neural network implementation of the random transitions here defined is discussed in (20, 21).

### C. Comparison between data and model simulation

We now turn to deploying this model in simulating the experimental paradigm analyzed previously. To qualitatively compare the model to experimental findings, we examine how the sequences generated by our model present grouping of items as measured by *p*_16_. In the model, the parameter *α* controls the relative strength of the list representation and thus higher levels of list grouping are achieved for higher values of *α*. We let *α* vary and with it we vary *β* according to 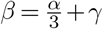 where *γ* is a constant that allows the contribution of session part of similarity to remain positive when list grouping controlled by *α* approaches zero. This is important as in the data we notice the two trends to be independent: even in FFR sessions that do not show any grouping induced by the IFR trial identity, words which were retrieved in a IFR trial had a higher probability to be retrieved. Using the described model, 6500 sessions of FFR were simulated. Each session first comprises the simulation of 16 IFR followed by FFR session, starting with a word previously recalled in IFR. All IFR sessions were simulated according to a non-hierarchical model where only *S*_*word*_ contributes to the similarity matrix, the matrix *S*_*tot*_ was computed and the FFR simulated for each of the 13 values of *α* linearly spanning the interval [0, 60].

We compute *p*_16_ for all sessions of FFR so generated and find that *p*_16_ monotonically increases with the value of *α*, Fig. 5a. This is an expected behavior since large values of *α* force structured recall. Similarly to experimental data the model shows a linear dependence of the number of recalled words as a function of *p*_16_, Fig. 5b (cfr. Fig. 1b). Intriguingly, the number of sequences of words recalled from the same list as a function of *p*_16_ shows a non-monotonic dependence, Fig. 5c (blue data), which we also observed in the data (orange data). For small values of *α*, and thus *p*_16_, the recall is unstructured and the number of sequences is roughly equal to the number of recalled words (see Fig. 1d). When *α* and, therefore, *p*_16_ increases the number of sequences increases since there is a mixture of two recall processes - random and structured (see Fig. 1e). For intermediate values of *p*_16_ the contribution of *S*_*word*_ and *S*_*list*_ to driving structured transitions are comparable and across lists transitions may still be triggered by structured transitions. As we further increase *α* the recall becomes very structured and the words from a single list are predominantly recalled before words from other lists are recalled (see Fig. 1c). Consequently the number of sequences becomes comparable, or even smaller than the number of presented lists. To further assess the validity of our model we compute the percentage of newly recalled words in FFR (the words that were not recalled in IFR). Fig. 5d shows that this steadily decreases with *p*_16_ for both the model (blue points) and the experimental data (orange points).

**Fig. 5.**
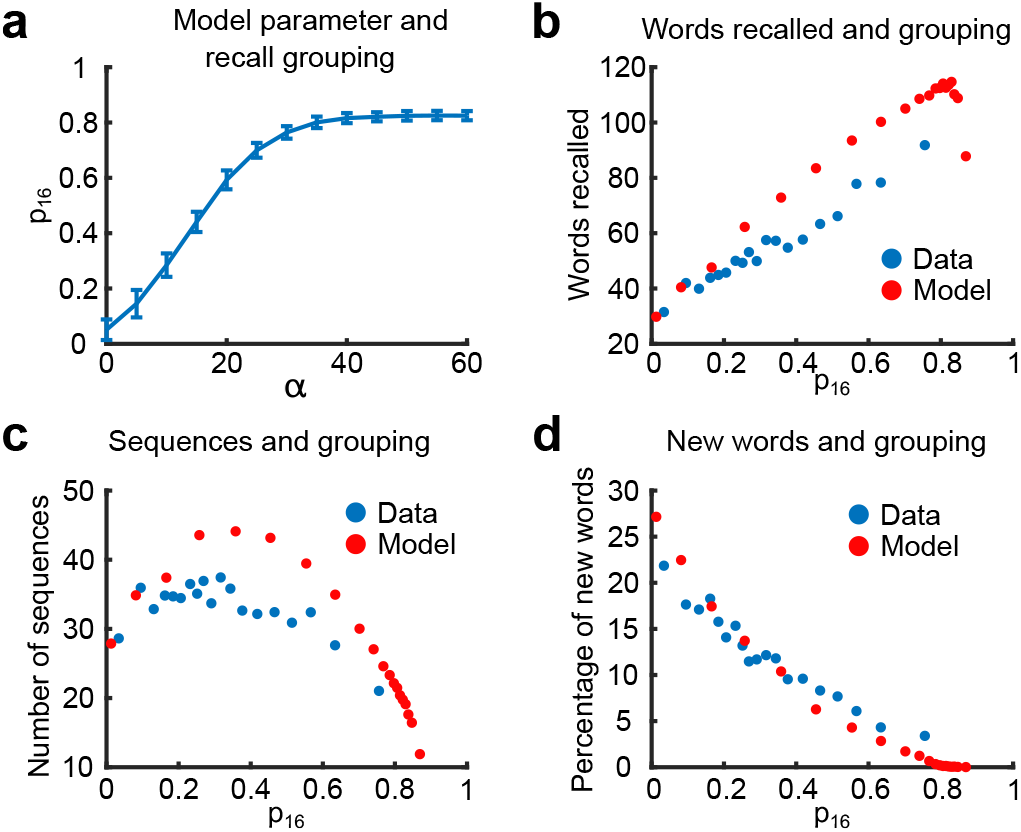
Hierarchical model results. **(a)** Value of the induced *p*16 in the model simulations as a function of *α*. Blue line is the average of the value of *p*16 with a confidence interval of 95%. **(b)** Scatter plot of the performance as a function of *p*16 both data and model. For each session the number of words retrieved in FFR is shown against the *p*16 score. **(c)** Scatter plot for the number of sequences retrieved as a function of *p*16 for each session both data and model. **(d** Scatter plot for the percentage of words not retrieved in the IFR but recalled in the FFR retrieved as a function of *p*16 for each session both data and model.

## Discussion

We studied the final free recall of sets of 256 unrelated words that were previously presented and recalled on the same day as 16 lists of 16 words each. We found that FFR trials exhibit various degrees of hierarchical organization: within-list chunking that spontaneously emerged in IFR, over-the-list organization induced by the presentation protocol, and finally list chunking for the very best participants (see Figs. 1 to 3 above). The dominant level, exhibited in the bulk of the data, was the tendency to recall subsequent words from the same list. This type of grouping strongly correlated with performance, cfr. Fig. 1b. When extrapolated to the limit of random recall, the performance dipped below the level of 30 words that closely matched our theoretical prediction for structure-less recall. The average performance was almost twice higher than this level, indicating a strong effect of information structure on memory retrieval. We also found that within-list chunks that emerged spontaneously in a limited number of trials in IFR (12) have a high probability to be recalled or omitted as single units during FFR trials as well. Taken together, our results strongly indicate that people tend to organize information to be remembered in a way that facilitate subsequent recall, even when information itself lacks any meaning, as in the case of free recall of random words. From a theoretical point of view we extended the model of associative memory recall (14) to take into account the hierarchical representation of information in FFR that we found in experimental data. More specifically, we added list and session context subnetworks including random and structured transitions. The effect of list and session context representations was limited to only words recalled in IFR and the strength of binding was a one-dimensional parameter in the model, *α*. The resulting model is compatible with proposed principles of sparse encoding and associative transitions. The simplicity of the model provides an interpretable description of the data as a function of strength parameter *α* providing insight to the processes involved in episodic memory storage and retrieval. The model is also easily generalizable to any number of hierarchical levels by adding additional layers of representations, similar to list representation.

In the current model the similarity matrix defining transitions consists of a weighted sum of similarity matrices describing the similarity in the different layers of hierarchy: words similarity, lists similarity and session context similarity. The binding between word similarities and lists similarities suggests the mechanisms for different observed phenomena. For example, both the non-monotonic dependency of the number of sequences and the constantly decaying number of newly recalled words in FFR are easily understood given the model (cfr. Fig. 5c and Fig. 5d). It can be argued that intrusions of words that were not presented during the day into FFR trials has a similar origin.

At the current level of realism, we propose to view the presented model as a platform for further development of realistic neural network models of information retrieval and other related types of cognitive tasks. Altogether we are able to interpret the complex structure of FFR data in terms of a model build on first principles (encoding and associativity). The capability of our model to shed light on multiple features of the dataset we analyze exploits the pivotal role of hierarchical representations in memory retrieval processes. We thus bridge between the cognitive ability of retrieval and the underlying neural dynamics pointing out, in simple terms, how individual differences in the ability of retrieving information may be interpreted by simple hierarchical properties of the encoded word representations.

## Materials and Methods

### Experimental methods

The data reported in this manuscript were collected in the lab of M. Kahana as part of the Penn Electrophysiology of Encoding and Retrieval Study (see (13) for details of the experiments). Here we analyzed the results from the 217 participants (age 17 - 30) who completed the first phase of the experiment, consisting of 7 experimental sessions. Participants were consented according the University of Pennsylvania’s IRB protocol and were compensated for their participation. Each session consisted of 16 lists of 16 words presented one at a time on a computer screen and lasted approximately 1.5 hours. Each study list was followed by an immediate free recall test. Words were drawn from a pool of 1638 words. For each list, there was a 1500 ms delay before the first word appeared on the screen. Each item was on the screen for 3000 ms, followed by jittered 800 - 1200 ms inter-stimulus interval (uniform distribution). After the last item in the list, there was a 1200 - 1400 ms jittered delay, after which participants were given 75 seconds to attempt to recall any of the just-presented items. In 4 out of 7 experimental sessions, following the immediate free recall test from the last list, participants were shown an instruction screen for final-free recall, informing them to recall all the items from the preceding lists in any order. After a 5 s delay, a tone sounded and a row of asterisks appeared. Subjects had 5 minutes to auditory recall any item from the preceding lists.

### Grouping measures

For each final-free recall trial we consider the ordered set of recalled words (*W)* defined as *w*_1_ → *w*_2_ →…→*w*_*n*_ where *n* is the number of words recalled in a given trial and *w*_1_ (*w*_2_, *…, w*_*n*_) denotes the input serial position during the day of the first (second,…,last) word recalled, which is the number between 1 and 256 (see Fig. 1a). We introduce the grouping measure (*p*), and assign the probability to each transition by assuming that the next word recalled is chosen from the same list as the currently recalled word with probability p and a random word is chosen with probability 1 - *p*. The probability for the whole sequence is computed as a product of individual transition probabilities. Formally, if *l*_*i*_ is the number of the list (from 1 to 16) from which word *w*_*i*_ was presented, the probability *P*_*i*_ of transition (*w*_*i*_ → *w*_*i*+1_) and the total logarithm probability of the whole sequence (log-likelihood) are

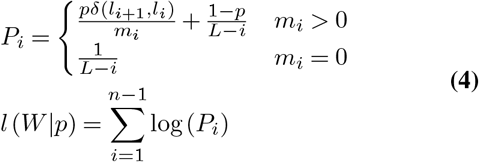

where *m*_*i*_ ∈ [0, *…*, 15] is the number of not yet recalled words from the list *l*_*i*_ ∈ [1, *…*, 16] and *L* = 256 is the total number of words presented during the day. The grouping measure *p*_16_ for the FFR trial is then obtained as the value of *p* that maximizes the likelihood of the sequence *l* (*W* |*p*).

### Theoretical model details

The model builds on the idea that words are represented as binary population vectors, Eq. (1). The full similarity matrix between two words is then given by Eq. (2). Given two words *W*_1_ and *W*_2_ the contributions of the different terms is given by

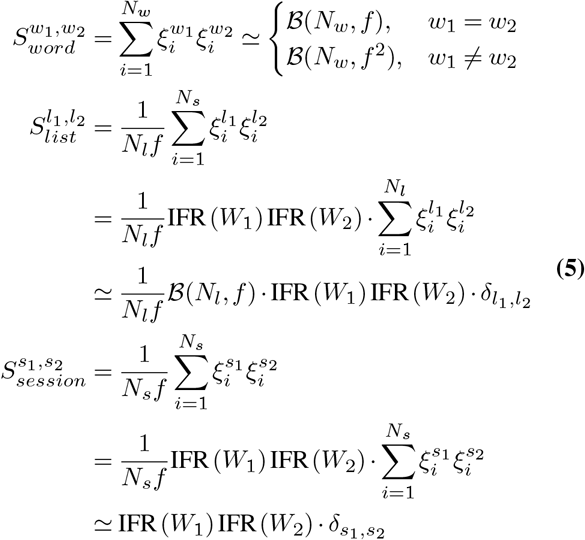

where *w*_1_, *w*_2_ index the word coding part of the population vector *ξ, l*_1_, *l*_2_ the list coding part, *s*_1_, *s*_2_ the session context coding part, while IFR(*W)* is an indicator function that word *W* was recalled in the IFR trial following the list presentation, i.e. IFR(*W)* = 1 was retrieved and 0 otherwise. To speedup simulations we neglected the correlations between elements in similarity matrices and approximate them by independent binomial random variables ℬ (*N*_*w*_, *f*^2^) for word similarities and ℬ (*N*_*l*_, *f)* for list similarities. We neglected similarities between different lists and different sessions and also assumed session similarities to be equal to each other. The list and session similarity matrices were normalized to have entries on the order of 1.

According to the associative principle, given an active word *W*_*k*_ the formal equation that defines the next word retrieved during structured recall is

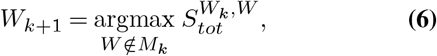

where *M*_1_ = {*W*_1_}, and *M*_*k*_ = {*W*_*k-*1_, *W*_*k*_}. Similarly for random transitions

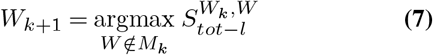

where *S*_*tot*-*l*_ is given by Eq. (3).

In the simulations we consider *N* = 300000 and *f* = 0.1, 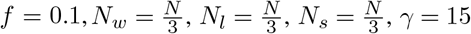. This value for *γ* was chosen to match the proportion of new words recalled during FFR on sessions with little over the list grouping (see Fig. 5d).

## ACKNOWLEDGEMENTS

This work is supported by the EU-H2020-FET 1564 and Foundation Adelis and EU - M-GATE 765549. We are grateful to M. Kahana for generously sharing the data obtained in his laboratory. The lab of Kahana is supported by NIH grant MH55687.

